# Computation-guided optimization of split protein systems

**DOI:** 10.1101/863530

**Authors:** Taylor B. Dolberg, Anthony T. Meger, Jonathan D. Boucher, William K. Corcoran, Elizabeth E. Schauer, Alexis N. Prybutok, Srivatsan Raman, Joshua N. Leonard

**Author notes:** These authors contributed equally to this work (Taylor B. Dolberg, Anthony T. Meger). **Contact Information**: Corresponding authors: Srivatsan Raman and Joshua N. Leonard.

## Abstract

Splitting bioactive proteins, such as enzymes or fluorescent reporters, into conditionally reconstituting fragments is a powerful strategy for building tools to study and control biochemical systems. However, split proteins often exhibit a high propensity to reconstitute even in the absence of the conditional trigger, which limits their utility. Current approaches for tuning reconstitution propensity are laborious, context-specific, or often ineffective. Here, we report a computational design-driven strategy that is grounded in fundamental protein biophysics and which guides the experimental evaluation of a focused, sparse set of mutants—which vary in the degree of interfacial destabilization while preserving features such as stability and catalytic activity—to identify an optimal functional window. We validate our method by solving two distinct split protein design challenges, generating both broad insights and new technology platforms. This method will streamline the generation and use of split protein systems for diverse applications.

## INTRODUCTION

Split proteins and conditional reconstitution systems are powerful tools for interrogating biology and controlling cell behavior.^1–4^ These systems work by splitting a protein into two fragments to disrupt the protein’s function. Each fragment is then fused to a partner domain such that the split protein is reconstituted, and its function is restored only when the partner domains interact. This modular strategy may be applied to diverse functional proteins to control bioluminescence^5, 6^, fluorescence^7^, proteolytic cleavage^8–10^ and transcription^11, 12^. As a result, conditionally-reconstituted split proteins have been employed in a variety of applications including probing and discovering new protein-protein interactions^13–16^, studying post-translational modifications^17^, imposing small molecule-regulated control over enzymatic activity^18, 19^, and rewiring cellular signaling^9, 20^.

Despite their utility in certain contexts, broader application of split protein systems is largely limited by the spontaneous reconstitution of fragments, resulting in high background activity (**Fig. 1a**). Splitting a protein tends to expose its hydrophobic core, creating highly unfavorable interactions between the core and solvent. Reconstitution is driven by a strong inherent preference to desolvate by recombining the fragments. Evaluating alternative splitting sites can vary reconstitution propensity, but this approach often only partially ameliorates the problem because changing splitting sites may not significantly affect underlying hydrophobic forces. Therefore, it is necessary to identify variants with a reconstitution propensity that precludes spontaneous reconstitution but enables reconstitution under desired conditions. Variants with a range of reconstitution propensities can be generated by random mutagenesis and screened for the desired property. However, high-throughput screening is not readily available for all split protein systems, and low-throughput clonal testing of variants can be laborious and suffer from inefficient exploration of sequence space. Even when screens generate improved variants, it may be difficult to interpret why only certain mutants were successful, and as a result, generalizable rules cannot be transferred to guide the tuning of new split protein systems. Furthermore, split protein systems tuned by mutagenesis exhibit performance characteristics determined by (and limited to) the conditions used in the initial screens, again posing a barrier to general applications.

**Fig. 1.**
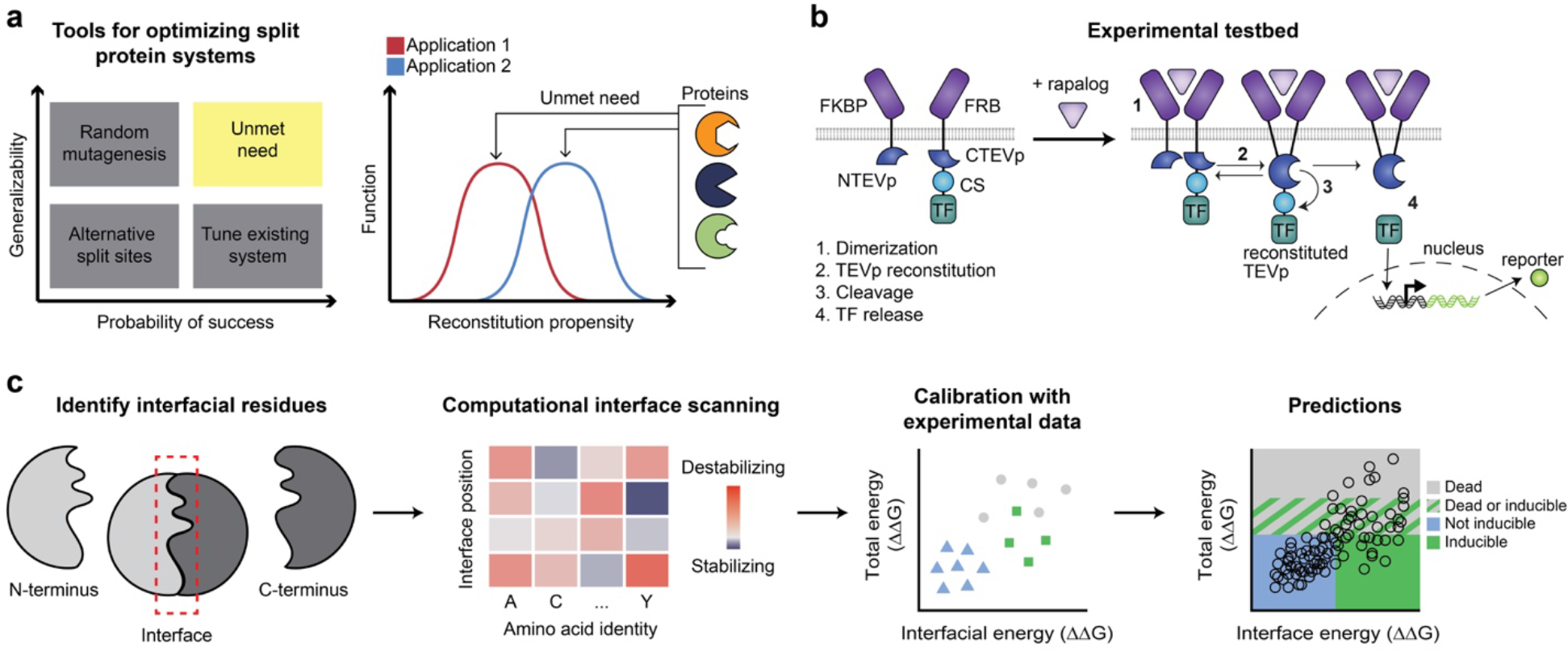
Design-driven strategy for tuning split protein systems. **a,** Current methods for optimizing split proteins are limited (left); an ideal tool would enable adapting split proteins for multiple applications, each of which may require distinct reconstitution propensities (right). **b,** This cartoon illustrates the experimental testbed used here; ligand binding-induced chain dimerization results in split TEVp reconstitution, trans-cleavage, and release of a previously sequestered transcription factor to drive reporter expression. **c,** <u>S</u>plit <u>P</u>rotein <u>O</u>ptimization by <u>R</u>econstitution <u>T</u>uning (SPORT) workflow: identify important residues at the split interface which are mutable, identify mutations that alter the total and interfacial energy, and use an application-specific model, trained on limited experiments, to identify those mutations that are predicted to yield a desired functional phenotype.

Here, we report a general strategy based on fundamental principles of protein biophysics for optimizing split protein systems which we term <u>**S**</u>plit <u>**P**</u>rotein <u>**O**</u>ptimization by <u>**R**</u>econstitution <u>**T**</u>uning, or SPORT. We use computational mutagenesis with the Rosetta macromolecular modeling suite to guide limited experimental screening and thereby discretely map the sequence-energy landscape of the split interface. This allows us to determine optimal interaction energies that maximize the performance of the split protein system. We demonstrate proof-of-concept by optimizing a split protease system for conditional reconstitution in two different contexts: membrane-embedded and cytosolic. Our approach generates simple design rules that may be extended to tune other split protein systems for distinct design goals and can be implemented by most research laboratories. This work demonstrates a new method for efficiently engineering split protein systems, which will streamline the generation and expand the use of split protein systems for diverse applications.

## RESULTS

### Formulation of the design challenge and strategy

As a first step toward developing a strategy that addresses the challenge of designing split proteins, we selected a model system based on the well-studied Tobacco Etch Virus protease (TEVp)—we sought to tune the reconstitution propensity of split TEVp. For this purpose, we modified a synthetic receptor system that we previously reported (Modular Extracellular Split Architecture, or MESA)^21^ to serve as a reporter of conditional split TEVp reconstitution. In this testbed, ligand binding-induced dimerization of a membrane receptor reconstitutes an intracellular split TEVp, which then autolytically liberates a sequestered transcription factor to drive reporter gene expression (**Fig. 1b**). Our initial evaluation demonstrated that the canonical split TEVp (split between residues 118/119)^22^ showed high propensity to reconstitute, resulting in high background from ligand-independent signaling (**Supplementary Fig. 1**). As the original screens used to identify this split were performed in a soluble rather than membrane-bound context, these data suggest that tethering split TEVp to a membrane may promote reconstitution. Furthermore, we determined that this problem is not limited to the canonical split site, as other TEVp partitioning also yielded poor performance (**Supplementary Fig. 2**). Given these observations, we formulated a design goal: rationally mutate split TEVp to optimize two key MESA performance characteristics—minimal reporter gene expression in the absence of ligand and a substantial fold increase in reporter expression upon ligand addition.

### Biophysical principles underlying SPORT

We developed SPORT, a computation-guided workflow to rationally design split protein interfaces to optimize reconstitution propensity (**Fig. 1c**). SPORT employs Rosetta, a state-of-the-art software package for protein design.^23^ Given a protein with a predetermined split site, our first step was to identify key interfacial residues to target for mutagenesis. Residues with large differences in solvent-accessible surface area (SASA)—when comparing intact protein and split fragments—were classified as buried residues. These buried residues are ideal targets for mutagenesis as they likely contribute substantially to the driving force for spontaneous reconstitution. For each buried residue, we performed a comprehensive, *in silico* mutational scan to evaluate the energy perturbation of all possible single-point mutations on the interaction energy across the split protein interface (ΔΔG_Interfacial_) and total stability of the mutated protein (ΔΔG_Total_) relative to the parent. The degree of disruption is a critical design consideration. Insufficient disruption may retain high background activity while excessive disruption may impair catalytic activity due to loss of overall protein stability. Therefore, the interface must be carefully tuned so that the driving force provided by ligand binding-induced dimerization promotes reconstitution. This “Goldilocks zone” likely differs for each individual protein and perhaps depends upon context, and this zone is difficult to define *a priori*. Therefore, our strategy was to identify the Goldilocks zone for a given protein by choosing mutations that span the range of ΔΔG values. We hypothesized that a limited test set of mutants would direct subsequent mutagenesis efforts by predicting desirable mutant combinations from a vast amount of sequence space. Each of these propositions was tested using experimental case studies.

### Validating SPORT by tuning a membrane-tethered split TEV protease

To investigate and validate SPORT, we applied our design workflow to the split TEVp MESA system. We first assessed the per-residue change in solvent accessible surface area (ΔSASA) between the intact form and split fragments (**Fig. 2a**). In total, 130 of the 218 residues showed increased SASA in the isolated fragments. We excluded from this set the catalytic triad and 27 residues lying within a 6 Å coordination sphere around the catalytic triad to avoid perturbing the catalytic function of reconstituted TEVp. Of the remaining 100 positions, we chose the 15 positions with the largest ΔSASA (9 in N-terminal and 6 in C-terminal halves of split TEVp) as candidates for mutagenesis. Next, we evaluated the energy perturbation of all possible single-point mutations (285 in total) at these positions using Rosetta. As expected, few mutations were predicted to increase stability, and a vast majority were destabilizing (**Fig. 2b**, **Supplementary Figure 3**). Many positions exhibited a variety of stabilizing, benign, and destabilizing point mutations that were consistent with structure. For instance, bulky sidechain substitutions (W, F, Y, R, K and H) at position 103 resulted in many steric clashes with neighboring residues (**Fig. 2b** right panel) and subsequently conferred large decreases in predicted stability.

**Fig. 2.**
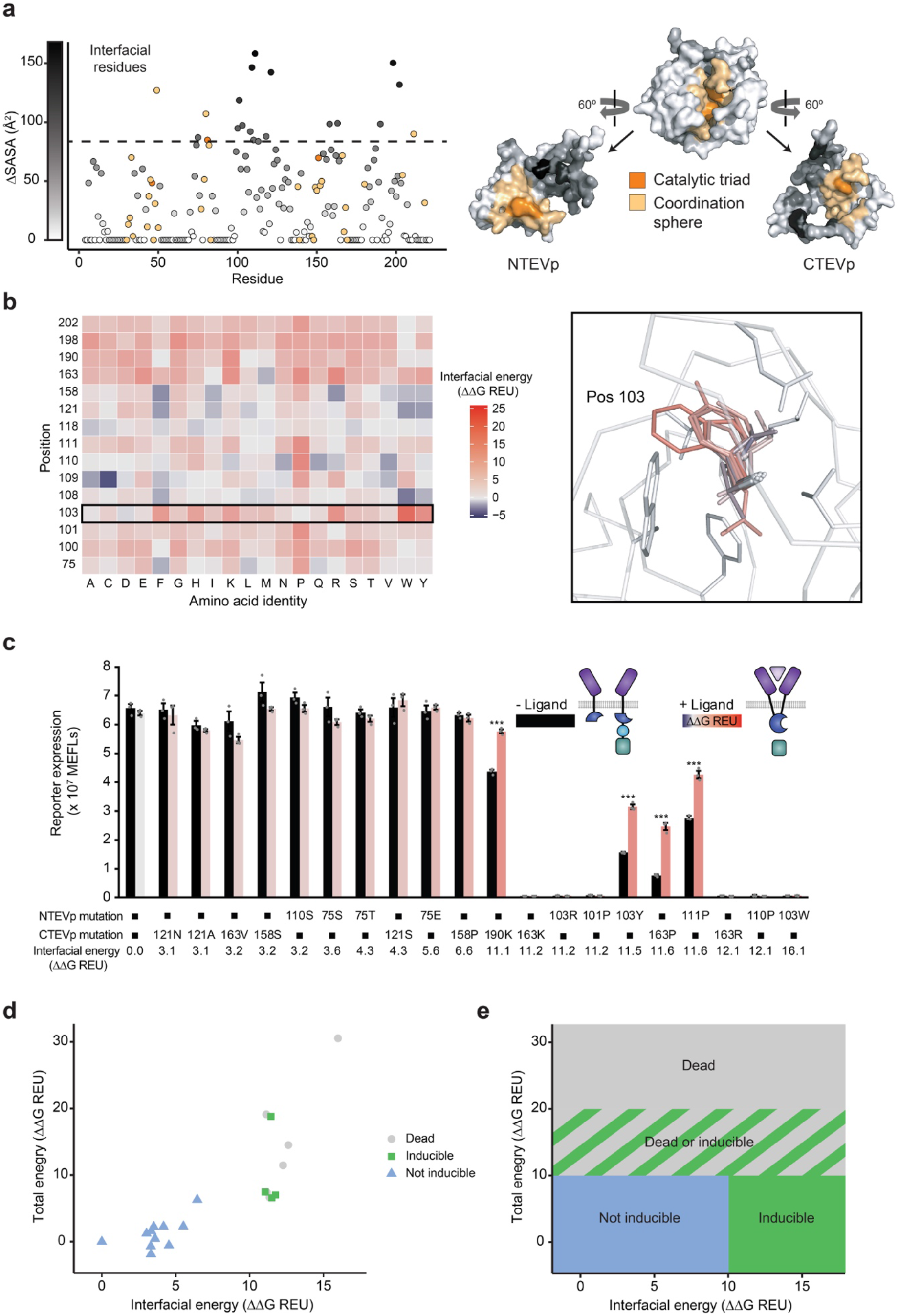
Computation guided method development and experimental analysis. **a,** Left, characterization of the solvent accessible surface area (SASA) of each residue of 118/119 split TEVp. Right, 3D depiction of 118/119 split TEVp, showing the catalytic triad (orange), coordination sphere (yellow), and ΔSASA (greyscale). **b,** Mutational scanning of high ΔSASA residues (left) and example of all possible mutations of residue 103 (right), with change in interfacial energy indicated by color. **c,** Experimental analysis of TEVp mutations predicted to span a range of interfacial energies. Error bars depict S.E.M. (*p ≤ 0.05, ***p ≤ 0.001). **d,** Experimental phenotypes observed in **c** were plotted on an energy landscape and annotated as indicated by color (reporter expression normalized to WT < 0.05 is “dead”, fold induction < 1.2 is “not inducible”, fold induction ≥ 1.2 is “inducible”). **e,** Proposed model for predicting zones of functional phenotypes based upon total and interfacial energy; the boundaries were proposed based upon observations using the initial 20 mutants tested in **c**.

Guided by these predictions, we next experimentally characterized 20 single-mutant split TEV variants that span a wide range of ΔΔG_Interfacial_ (3.1 to 16.1 Rosetta Energy Units, or REU) and ΔΔG_Total_ (−1.9 to 30 REU) energies (**Fig. 2c**). We observed high background signaling (i.e., reporter expression) for disruptions up to ΔΔG_Interfacial_ ~6.6 REU, which suggested that destabilization was insufficient. However, four out of ten variants with ΔΔG_Interfacial_ >10 REU exhibited reduced background signaling and substantial ligand-induced activation (**Fig. 2c**). The remaining six were completely inactive (or “dead”); they induced no signaling under any conditions. Energy-based partitioning across different phenotypes (inducible, not inducible, and dead) became evident when comparing ΔΔG_Interfacial_ and ΔΔG_Total_ of all 20 single mutant split TEV variants (**Fig. 2d**). Variants with high background activity due to insufficient destabilization fell in the region where ΔΔG_Interfacial_ < 10 and ΔΔG_Total_ < 10 REU. Variants with the dead phenotype had ΔΔG_Interfacial_ > 10 and ΔΔG_Total_ > 10 REU; since these mutants were expressed, as confirmed by Western blot (**Supplementary Fig. 4**), the lack of signaling suggested that these mutations directly preclude reconstitution. Most of the inducible variants (three out of four) were observed in the energy window where ΔΔG_Interfacial_ > 10 and ΔΔG_Total_ < 10 REU, which may represent the Goldilocks zone we hypothesized to exist. An additional region contained a mixture of inducible and dead phenotypes. By inspection of these results, we then proposed a model for broadly classifying experimental phenotypes based on energy partitions (**Fig. 2e**).

### SPORT predicts outcomes of combining mutations

We next evaluated whether our proposed classifier model—developed based upon observations with single mutants—could predict the phenotypes of combined NTEVp and CTEVp mutations, including both double (two mutations on one chain) and paired (one mutation on each chain) mutants derived by combining the initial 14 single non-dead mutations tested (**Fig. 3a**). Of the 67 possible double and paired mutants tested, 28 were predicted to be inducible. We experimentally tested 14 of these and found that 10 exhibited inducible signaling as predicted, one was dead, and three were not inducible; this yields an observed accuracy of 0.71 for inducible predictions (**Fig. 3b,c**). Interestingly, three of the prediction failures fell at the low end of the range of predicted changes in interfacial energy, suggesting potential opportunities for refining the classifier model. We also noted an interesting trend in our ΔΔG (total and interfacial) calculations—for the sixty-seven mutants tested, the calculated ΔΔG for the double and paired mutants were nearly identical to the sums of the ΔΔG calculated for the associated single mutants (**Supplementary Fig. 5**). Thus, for subsequent analyses of combined mutants, we simply added the effects of single mutants in our calculations.

**Fig. 3.**
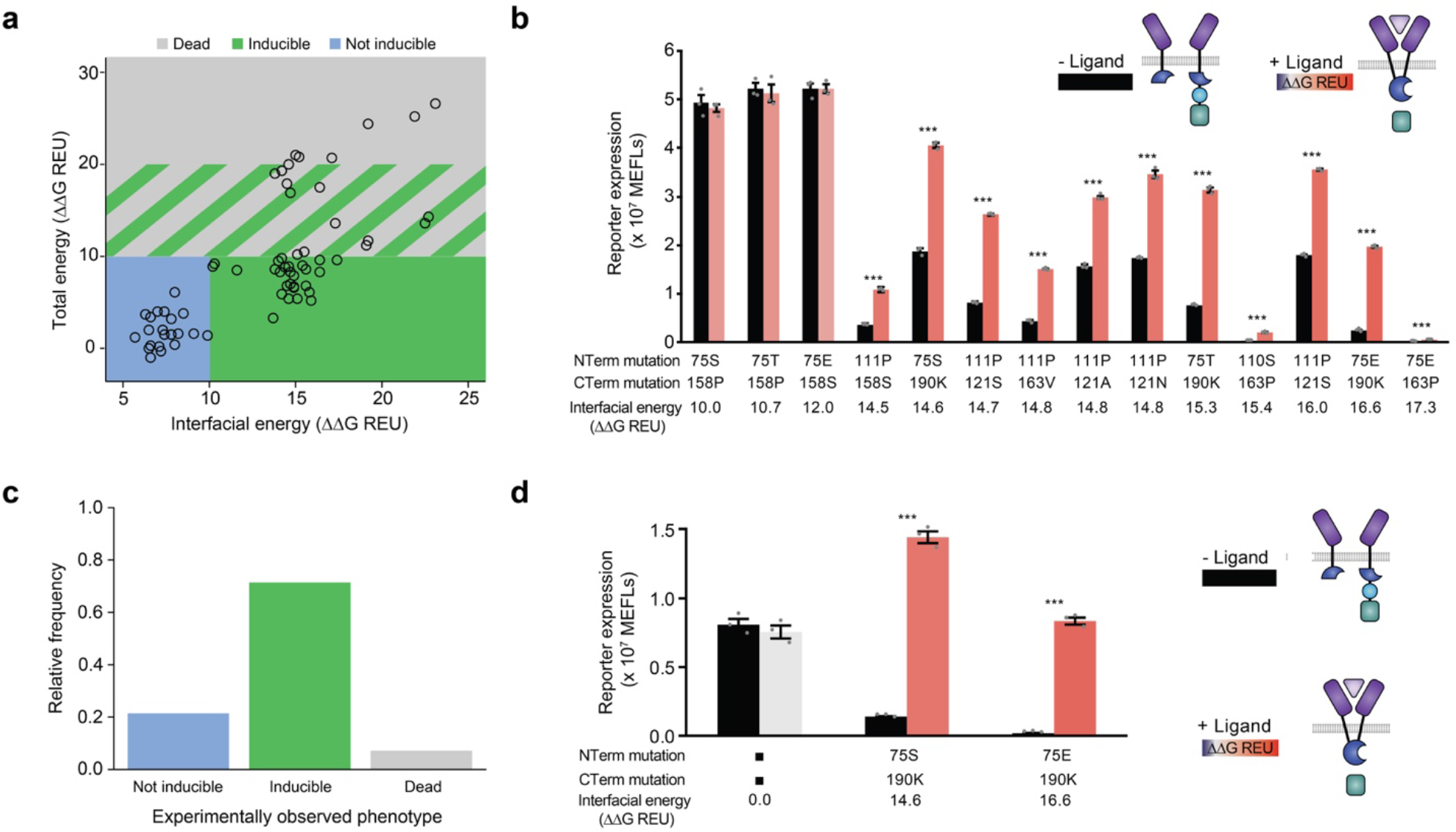
Evaluation of model-predicted phenotypes for combined mutations. **a,** Computed energies and predicted phenotypes based on the classifier model—proposed in **Fig. 2e**—of all possible double and paired mutants constructed by combinatorial sampling of the initial single mutants tested (omitting dead mutations) in **Fig. 2c**. **b,** Experimental evaluation of selected mutants predicted to be inducible. **c,** Experimentally observed phenotypes for the fourteen mutants predicted to be inducible (from **b**), showing that the model predicts inducibility at a fairly high rate (10/14). **d,** Normalizing protein expression levels improves performance (fold induction) of selected mutants (from **b**), whereas WT function is not changed. Normalization was achieved using Western Blot analysis (**Supplementary Fig. L4)** to adjust DNA doses transfected (per well, N-terminal chains: 0.4 ng WT, 1 ng 75S, 1.4 ng 75E; C-terminal chains: 5 ng WT, 12 ng 190K). Error bars depict S.E.M. (*p ≤ 0.05, ***p ≤ 0.001).

We next investigated how variations in expression level might impact the inducibility of the mutants. We used Western blot analysis to normalize and vary chain expression levels by adjusting DNA doses (**Supplementary Fig. 6**). Notably, these constructs remained inducible across the entire range of expression levels tested, suggesting that the biophysical mechanism of optimized split protein reconstitution is robust to variations in the expression levels and ratio of the membrane-bound split TEVp fragments. However, we observed that the performance of these constructs (i.e., fold induction of signaling upon ligand addition) could be substantially improved through tuning expression such that protein levels of each fragment are comparable (**Fig. 3d**, **Supplementary Fig. 7**). Taken together, these results suggest that a classifier calibrated with a limited set of experimental observations spanning the full range of ΔΔG can predict function of new mutants with high accuracy in a manner that is independent of expression level of the construct.

### SPORT predicts phenotypes of novel mutations and combinations

All mutations previously tested were derived from predictions based on our computational method. Next, we wanted to test a broad set of mutants (outside the calibration set) to investigate the accuracy of our classification scheme. Therefore, we experimentally characterized additional single mutants (not included in the original calibration set) and all combinations of paired mutations derived from both the original and this expanded mutation set (omitting dead constructs); mutants were selected to explore the boundaries of the energy landscape classifier model and were expected to reflect a wide range of induced and uninduced reporter expression levels and ratios (**Figs. 2e**, **3a**). We also sought to investigate whether the phenotypic partitioning demonstrated for inducibility (**Fig. 3b,c**) is extensible to the other phenotype classes (i.e., dead and not inducible). This expanded set paired 10 N-terminal mutations with 16 C-terminal mutations. In general, we observed that variants with larger ΔΔG_Interfacial_ energies had lower reporter expression levels in both the background (OFF) and ligand-induced (ON) states (**Fig. 4a**, left and middle panels). Thus, the cost of lowering the OFF state is to also lower the ON state, but these reductions are not always proportional. This is evident by the diversity of calculated fold inductions (**Fig. 4a**, right panel). Only one variant, H75P/W198E, exhibited a significant decrease in the OFF state and an increase in the ON state relative to wild type (WT). However, the variants with highest fold inductions, such as H75S/I163P (17.3 fold induction) and H75T/I163P (9.92 fold induction), exhibited significantly lower OFF and ON states than did the WT, reflecting a tradeoff between desirable performance characteristics. Overall, we observed moderate agreement between the actual and predicted phenotypes for these novel variants and combinations (**Fig. 4b**). The classifier model was most accurate for predicting the not inducible phenotype (25 of 29, 86%). Many inducible phenotype predictions were also confirmed (31 of 52, 60%). This success rate is impressive given that phenotypic classification boundaries were set roughly based upon the sparse calibration set (**Fig. 2e**). Interestingly, this analysis also indicates that the energy landscape calculated by SPORT correlates with each phenotype to differing degrees.

**Fig. 4.**
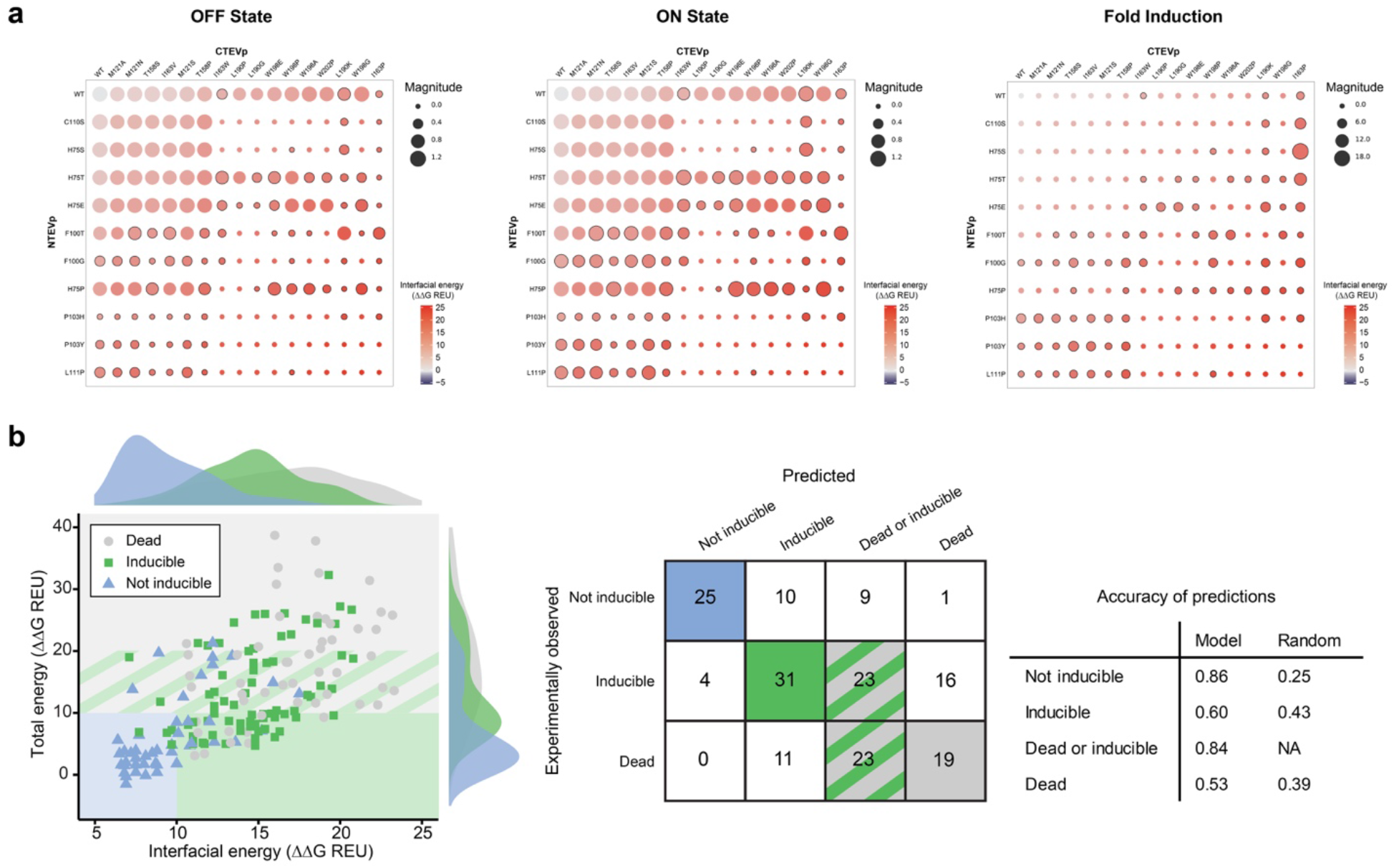
Evaluation of model-predicted phenotypes for novel mutations and combinations. **a,** For each experimentally characterized construct, reporter output was quantified in the absence of ligand (OFF state) and following ligand addition (ON state), and the fold induction was calculated. Calculated interfacial energy for each construct is indicated by circle *color*, magnitude of reporter expression (or fold-induction) is indicated by circle *size*, and constructs with a fold induction ≥ 1.2 are denoted with a black border. Single mutants observed to be dead (**Fig. 2**) were not carried forward to this analysis. **b,** Left, experimentally observed phenotypes (data point color) were mapped onto the proposed classifier model from **Fig. 2e**, with observed frequency distributions shown as histograms. Right, evaluation of model prediction accuracy compared to random assignment of phenotypes.

In order to gain additional insight into how choice of calibration set and sample size may impact the accuracy of SPORT predictions, we performed retrospective bootstrapping analysis of the data presented in **Figure 4** (see **Supplementary Note 2** for full details). Experimental data were stratified by the energy landscape and partitioned randomly into calibration and prediction subsets. Logistic regression modeling was applied to evaluate the accuracy of classification under various conditions (**Supplementary Fig. 8**). We observed robust prediction accuracy using multiple unique calibration sets and using sample sizes ranging from 4 to 28. This outcome suggests that our ability to generate a general classification model was not dependent upon the specific calibration data we used in our initial characterization experiment (**Fig. 2**), and that a relatively small set of calibration data drawn from a distribution like that included in **Figure 4** would be sufficient to generate a general classification model.

### Extension of SPORT predictions to new design goals

A major limitation to current approaches for employing split proteins is that often a variant selected to perform well in one context fails in a different context. To investigate whether the SPORT design method is generalizable beyond our initial model system, we developed a distinct model system. This new system employs split TEVp in a soluble form, where we hypothesized that a different reconstitution propensity would be required compared to the membrane-bound model system. To generate such a soluble test system (**Fig. 5a**), ligand binding domains were fused to split TEVp domains along with a soluble transcription factor flanked by protease cleavage sites and nuclear export signals (NES); thus, TEVp-mediated cleavage removes the NES from the transcription factor to enable nuclear localization and reporter expression. We first developed and tested a panel of soluble transcription factors that could implement this mechanism. This evaluation included varying the number of NES elements, their placement at N and/or C terminus of the transcription factor, and the P1’ residue of the TEVp cleavage sequence which governs cleavage kinetics^24^ (**Fig. 5b**). Several soluble transcription factor constructs exhibited the desired phenotype of low signaling in the absence of TEVp and high signaling when co-expressed with full TEVp; construct TF10 was selected for evaluating split TEVp variants.

**Fig. 5.**
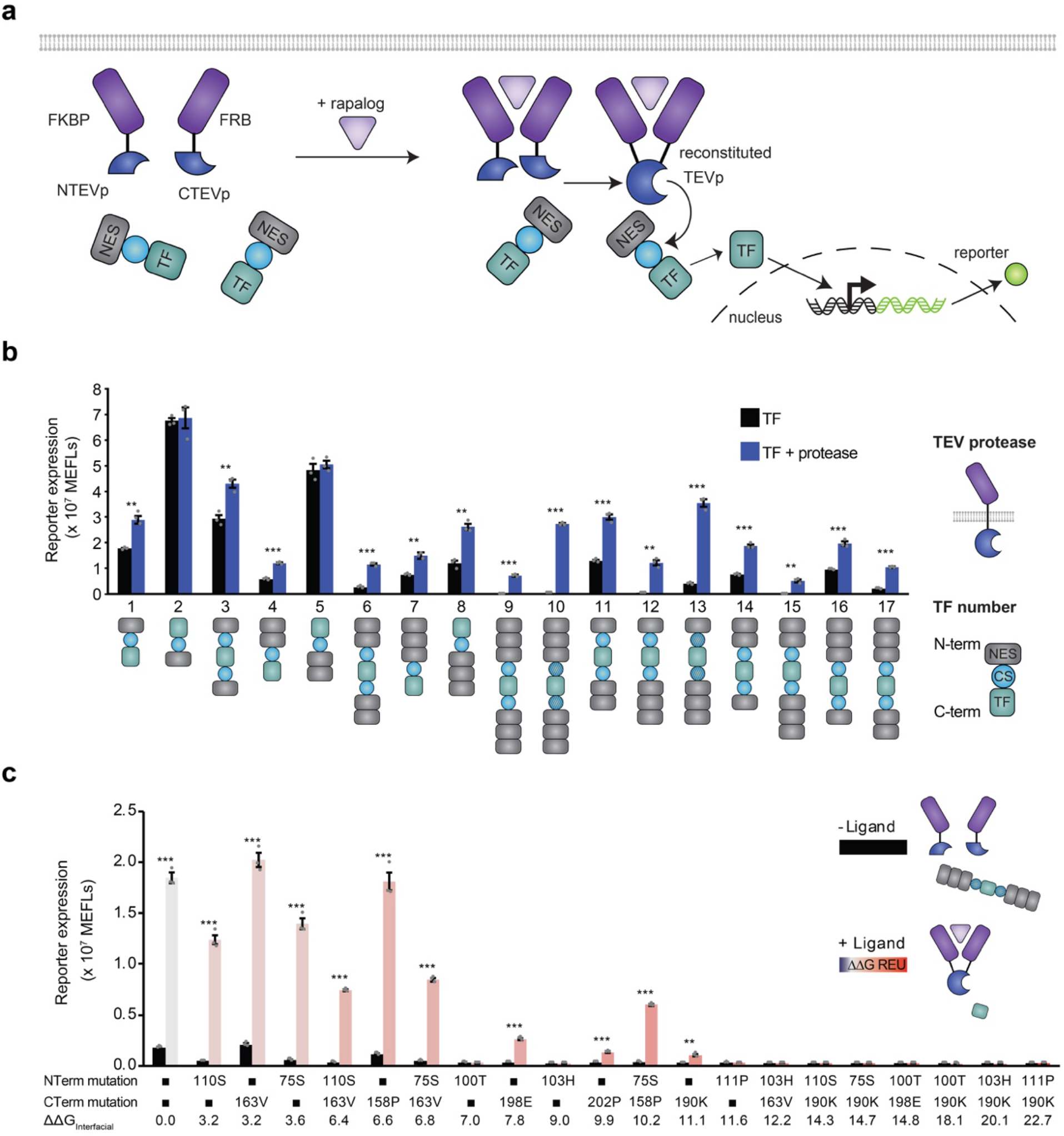
Model guided design of a new split TEVp application in soluble context. **a,** This cartoon illustrates the soluble split TEVp testbed. Ligand-binding-induced dimerization mediates reconstitution of split TEVp, which then cleaves one or more nuclear export sequence (NES) elements from a soluble transcription factor, leading to nuclear import and reporter expression. **b,** Developing the testbed by evaluating engineered transcription factors (TF) for consistency with the mechanism proposed in **a**; shaded cleavage sequence (CS) domains indicate a G residue in the P1’ position, unshaded CS domains indicate a M residue in this position.^24^ **c,** Experimental analysis of single and paired mutants sampling a range of interfacial energies (indicated by color and labeled), employing TF10 from **b**. Error bars depict S.E.M. (*p ≤ 0.05, ***p ≤ 0.001).

Using our soluble split TEVp test system, we evaluated a panel of 10 single TEVp mutants and 10 paired TEVp mutants spanning a range of interfacial energies **(Fig. 5c**). The construct based upon WT split TEVp exhibited a substantial fold induction, which is consistent with the fact that this split protein was identified by screens performed in the soluble phase^22^. However, the WT construct also yielded high background signaling in the absence of ligand, indicating an opportunity to improve performance. For modest increases in ΔΔG_Interfacial_ (~0–3.6 REU), background signaling persisted. For intermediate increases in ΔΔG_Interfacial_ (~6-10 REU), a mixture of phenotypes was observed, including dead constructs and those with both substantially reduced background signaling and substantial fold inductions. For large increase in ΔΔG_Interfacial_ > ~10 REU, constructs were generally weakly-inducible or dead. Thus, a focused evaluation of 20 design variants, guided by SPORT, yielded ~5 variants exhibiting improved performance compared to the WT construct. Altogether, these observations demonstrate that the SPORT method and associated energy landscape concept can be employed to efficiently solve distinct split protein optimization challenges.

## DISCUSSION

In this study, we developed and validated what is—to our knowledge—the first computational strategy for tuning split protein reconstitution propensity. Although the split TEVp MESA used as our first model system would have been deemed infeasible using standard evaluations of split proteins (**Supplementary Figs. 1-2**), application of SPORT to tune this system yielded multiple high-performing new synthetic receptor scaffolds (**Fig. 3d**). We show that unlike the classical MESA receptors we have characterized in prior work^21, 25, 26^, split TEVp MESA tuned by SPORT exhibit excellent performance characteristics (i.e., low-background and high fold-induction) in a manner that is robust to variations in both biosensor expression level and the ratio at which biosensor chains are expressed (**Supplementary Figure 7)**. This property is of great practical utility, as it precludes the need to carefully tune the implementation of each biosensor.

Several important insights emerged from this study. First, our approach demonstrated that testing a sparse set of mutants along the energy landscape is an effective strategy to choose optimal interfacial energies to promote conditional reconstitution. Second, multiple point mutations with similar energies exhibited similar performance, which suggests reconstitution propensity depends on the energy of destabilization but is agnostic to specific mutations. Third, the concept of a Goldilocks zone is likely generalizable to different proteins and application contexts, but the optimal energy window may have to be adjusted on a case-by-case basis. We find that membrane-bound split proteins must be destabilized to a greater degree than soluble split proteins in order to avoid spontaneous reconstitution. Altogether, these results show that split protein systems can be engineered based on fundamental principles of protein biophysics, which obviates the need for exhaustive screening and generates rules applicable to new candidate proteins.

There are several interesting opportunities for extending and improving SPORT in future work. First, although our analysis showed that SPORT can be used to identify mutations that confer specific energy changes, it does not yet enable *a priori* prediction of where the Goldilocks zone will fall for new applications. It is possible that subsequent analysis of many case studies could identify trends that enable such predictions and thus harness SPORT to further focus experimental investigations. An additional opportunity is pairing SPORT with a multiparameteric optimization framework for exploring pareto-optimal tradeoffs between performance characteristics; for example, in our model system, there seems to be such a tradeoff between low background in the ligand-free state and high signaling in the ligand-induced state. Finally, the SPORT algorithm itself may be refined to better avoid false positives (e.g., dead mutants that share a region of the energy landscape with inducible variants). Altogether, our findings suggest many opportunities for expanding the utility of split proteins for many new applications and highlight the impact of SPORT-guided development of novel biochemical and synthetic biology tools.

## METHODS

### General DNA assembly

Plasmid cloning was performed using standard molecular biology techniques of PCR and restriction enzyme cloning with Phusion DNA Polymerase (NEB), restriction enzymes (NEB; Thermo Fisher), T4 DNA Ligase (NEB), and Antarctic Phosphatase (NEB). Development of the tTA-responsive YFP reporter plasmid was described previously^21^. Plasmids were transformed into chemically competent TOP10 *E. coli* (Thermo Fisher) and grown at 37°C. Plasmid maps are provided as GenBank files (**Supplementary Data 1**).

### Plasmid preparation

Plasmid DNA used for transfection was prepared using the PEG precipitation method, which was previously described in detail.^27^

### Cell culture

HEK293FT cells (Life Technologies/Thermo) were maintained at 37°C incubator and 5% CO_2_. Cells were cultured in DMEM (Gibco 31600-091) with 10% FBS, 6 mM L-glutamine (2 mM from Gibco 31600-091 and 4 mM from additional Gibco 25030-081), penicillin (100 U/µL), and streptomycin (100 µg/mL) (Gibco 15140122).

### Transfection

Transfections were performed in 24 well plates seeded at 1.5 × 10^5^ cell in 0.5 mL of DMEM media. At 6-8 hours post-seeding, cells were transfected using calcium phosphate method with a total DNA content of 1-2 ug DNA per mL of media, using DNA prepared by PEG precipitation. All experiments included blue fluorescent protein (BFP) as a control to assess transfection efficiency. The exact DNA amounts added to the mix per well are as follows, unless otherwise stated in figure captions: 25 ng of each TEVp chain, 200 ng of BFP control, 200 ng of YFP reporter plasmid, and 150 ng of pcDNA plasmid. This mixture was added dropwise to an equal-volume solution of 2x HEPES-Buffered Saline (280 mM NaCl, 0.5 M HEPES, 1.5 mM Na_2_HPO_4_) and gently pipetted up and down four times. After 2.5 minutes, the solution was mixed vigorously by pipetting ten times and 100 μL of this mixture was added dropwise to each well of the plated cells, and the plates were swirled gently. For functional experiments, 12 hours post-transfection, media containing 0.1 μM rapamycin analog (Takara AP21967) or 0.1% ethanol as a control was added to cells. At 24-30 hours post-media change, cells were harvested for flow cytometry with Trypsin-EDTA, which was then quenched with medium, and the resulting cell solution was added to at least 2 volumes of FACS buffer (PBS pH 7.4 with 2–5 mM EDTA and 0.1% BSA). Cells were spun at 150 × g for 5 min, FACS buffer was decanted, and fresh FACS buffer was added. All experiments were performed in biological triplicate.

### Flow Cytometry

Approximately 10^4^ live cells from each transfected well of the 24-well plate were analyzed using a BD LSR Fortessa Special Order Research Product (Robert H. Lurie Cancer Center Flow Cytometry Core) running FACSDiva software. Samples were analyzed using FlowJo v10 software (FlowJo, LLC). The HEK293FT cell population was identified by FSC-A vs. SSC-A gating, and singlets were identified by FSC-A vs. FSC-H gating. A control sample of cells—generated by transfecting cells with a mass of pcDNA (empty vector) equivalent to the mass of DNA used in other samples in the experiment—was used to distinguish transfected and non-transfected cells. For the single-cell subpopulation of the pcDNA-only sample, a gate was made to identify cells that were positive for the constitutively driven blue fluorescent protein (BFP) used as a transfection control in other samples such that the gate included no more than 1% of the non-fluorescent cells. The mean fluorescence intensity (MFI) of the single-cell transfected population was calculated and exported for further analysis. BD LSR Fortessa settings used were as follows: BFP was collected in the Pacific Blue channel (405 nm excitation, 450/50 nm filer) and EYFP was collected in the FITC channel (488 nm excitation, 505 LP and 530/30 nm filter). To quantify reporter expression, the FITC channel MFI was averaged across three biological replicates. Cell autofluorescence was subtracted and MFI was converted to Mean Equivalents of Fluorescein (MEFLs) using the coefficient determined by the calibration curve of UltraRainbow Calibration Particles (Spherotech URCP-100-2H) run in each individual experiment. Standard error was propagated through all calculations.

### Western Blotting

Western blots were performed to evaluate protein expression and normalize total expression of each TEVp chain. A 3X-FLAG tagged NanoLuciferase as a normalization control, and images were analyzed using ImageJ software. A detailed western blot protocol was previously described.^27^

### Solvent-accessible Surface Area

The structure of TEVp was obtained from the Research Crystallography for Structural Bioinformatics (RCSB) PDB (ID code: 1LVM). Per-residue solvent-accessible surface areas (SASA) were computed using GROMACS v2018.1, which utilizes the double cubic lattice method (DCLM) described by Eisenhaber et al.^28^ The change in solvent accessible area was computed as

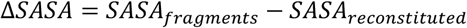

where structures of the N and C-terminal fragments were isolated from the crystal structure.

### Computational Interface Scanning

All modeling calculations were performed using the *Rosetta* molecular modeling suite v3.9. Single-point mutants were generated using the standard Relax application, which enables local conformational sampling to minimize energy (**Supplementary Note 1** includes full details). The total energy (ΔG_Total_) of each mutant was computed as the average of 100 relaxed models. The energy perturbation to total energy was computed as

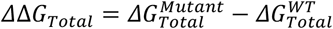

The Rosetta Scripts application with the InterfaceAnalyzeMover was applied to each relaxed model to compute the average residue-residue interaction energies between the N- and C-terminal fragments (**Supplementary Note 1** includes the full details). The interfacial energy was computed as the pair-wise sum of all short-range interaction energies as shown by

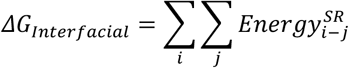

where i and j denote the sets of residues within each fragment. The energy perturbation of each mutation to the interfacial energy was then computed as

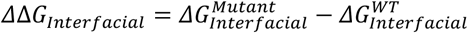

### Phenotype Classifier

Experimentally characterized variants were assigned class labels (not-inducible, inducible and dead) based on reporter expression levels in the ligand-absent and ligand-induced states. Variants with ≥1.2 fold higher reporter expression in the ligand-induced state relative to the ligand-absent state were labeled as inducible. For variants with expression levels <5% of wild-type (WT) sequence in the ligand-induced state and <1.2 fold activation were classified as functionally dead. The remaining variants were assigned the not-inducible class label.

## STATISTICAL ANALYSIS

Statistical details for each experiment are included the figure legends. The data shown reflect the mean across these biological replicates of the mean fluorescence intensity (MFI) of approximately 2,000–3,000 single, transfected cells. Error bars represent the SEM (standard error of the mean). For statistical analyses, two-tailed Student’s t-tests were used to evaluate whether a significant difference exists between two groups of samples, and the reported comparisons meet the two requirements of this test: (1) the values compared are expected to be derived from a normal distribution, and (2) the variance of each group is expected to be comparable to that of the comparison group since the same transfection methodologies and data collection methods were used for all samples that were compared. A p value of ≤ 0.05 was considered to be statistically significant.

## DATA AVAILABILITY

Data reported in composite figures (**Figs. 2a,b,d, 3a, 4b, Supplementary Fig. 4**) are included as **Source Data**.

## CODE AVAILABILITY

Rosetta details and script are provided in **Supplementary Note 1**.

## ACKNOWLEDGEMENTS

This work was supported in part by the National Institute of Biomedical Imaging and Bioengineering of the NIH under Award Number 1R01EB026510 (JNL); the Northwestern University Flow Cytometry Core Facility supported by Cancer Center Support Grant (NCI 5P30CA060553); T.B.D was supported by the Department of Defense (DoD) through the National Defense Science & Engineering Graduate Fellowship (NDSEG). This work is also supported in part by the Great Lakes Bioenergy Research Center, U. S. Department of Energy, Office of Science, Office of Biological and Environmental Research under Award Number DE-SC0018409 (S.R and A.T.M). The content is solely the responsibility of the authors and does not necessarily represent the official views of the NIH, Department of Defense, Department of Energy or other federal agencies.

## AUTHOR CONTRIBUTIONS

T.B.D., A.T.M., S.R, and J.N.L conceptualized the project. T.B.D., J.D.B, W.K.C., E.E.S, created reagents, designed and performed experiments, and analyzed the data. A.N.P. assisted in analyzing and visualizing the data. A.T.M., developed the computational model and code. T.B.D., A.T.M., S.R., and J.N.L. drafted the manuscript, T.B.D., A.T.M., and A.N.P. created the figures. J.N.L. and S.R. supervised the work. All authors edited and approved the final manuscript.

