## Supplementary Information for "Computation-guided optimization of split protein systems"

Taylor B. Dolberg[1,2,9], Anthony T. Meger[3,4,9], Jonathan D. Boucher[2,5], William K. Corcoran[2,5], Elizabeth E. Schauer[1,2], Alexis N. Prybutok[1,2], Srivatsan Raman[3,4,8,*], Joshua N. Leonard[1,2,5,6,7,*]

[1]Department of Chemical and Biological Engineering, Northwestern University, Evanston, Illinois 60208, United States

[2]Center for Synthetic Biology, Northwestern University, Evanston, Illinois 60208, United States

[3]Department of Biochemistry, University of Wisconsin-Madison, Madison, Wisconsin 53706, United States

[4]Great Lakes Bioenergy Research Center, University of Wisconsin-Madison, Madison, Wisconsin 53706, United States

[5]Interdisciplinary Biological Sciences Graduate Program, Northwestern University, Evanston, Illinois 60208, United States

[6]Chemistry of Life Processes Institute, Northwestern University, Evanston, Illinois 60208, United States

[7]Member, Robert H. Lurie Comprehensive Cancer Center, Northwestern University, Evanston, Illinois 60208, United States

[8]Department of Bacteriology, University of Wisconsin-Madison, Madison, Wisconsin 53706, United States

[9]These authors contributed equally to this work

**Contact Information**

**Contents:**

Supplementary Figures 1–8

Supplementary Table 1[†]

Supplementary Notes 1-2

[†] The contents of this table are provided separately as a Microsoft Excel Worksheet; the legend is provided here.

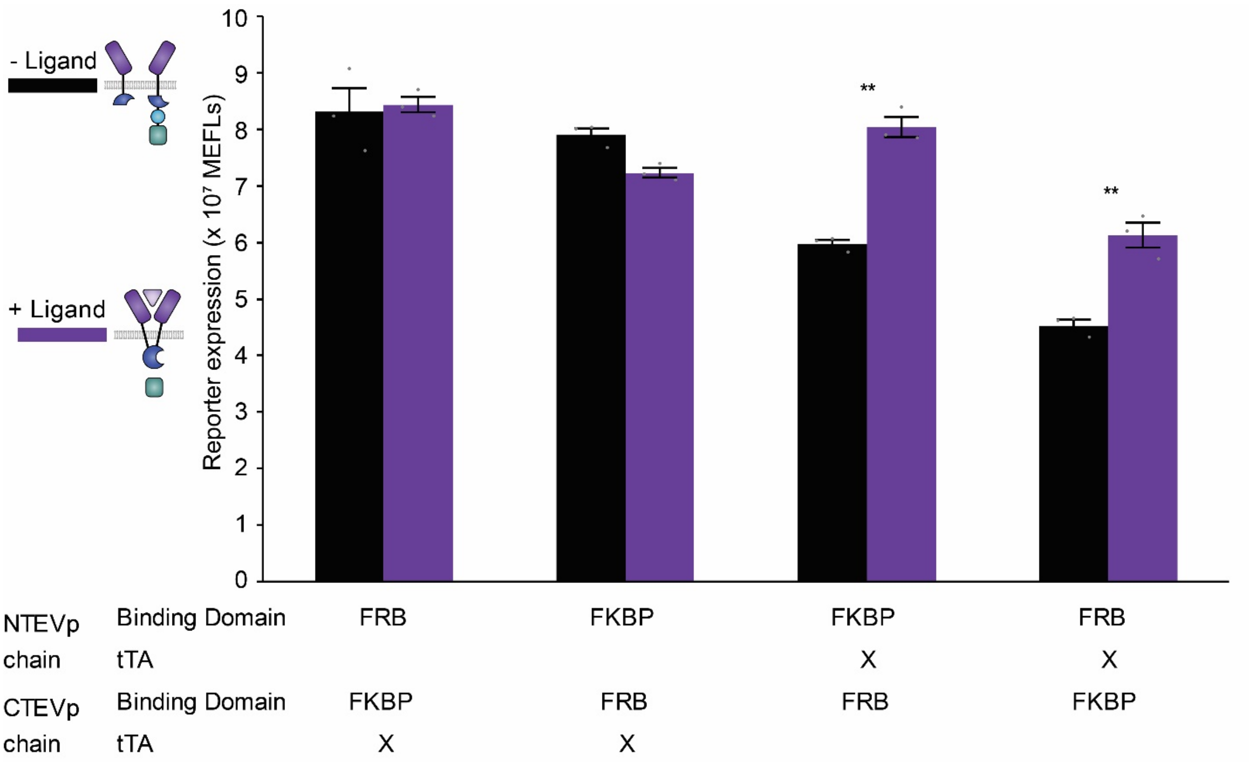


**Supplementary Fig. 1 Comparative evaluation of architectures for the membrane-bound model system.** Experimental analysis of various architectures of canonical 118/119 split TEVp MESA system, swapping the rapamycin binding domains (FRB, FKBP), TEVp fragment (NTEVp, CTEVp), and which chain contains the tTA. Reporter expression driven by reconstituted TEVp cleavage of tTA measured with flow cytometry. All four architectures were observed to have very high ligand independent signaling, suggesting that the 118/119 split of TEVp reconstitutes too easily without the addition of ligand, regardless of rapamycin binding domain and tTA chain architecture. The second architecture was carried forward for further analysis. Error bars depict S.E.M. (**P ≤ 0.01).

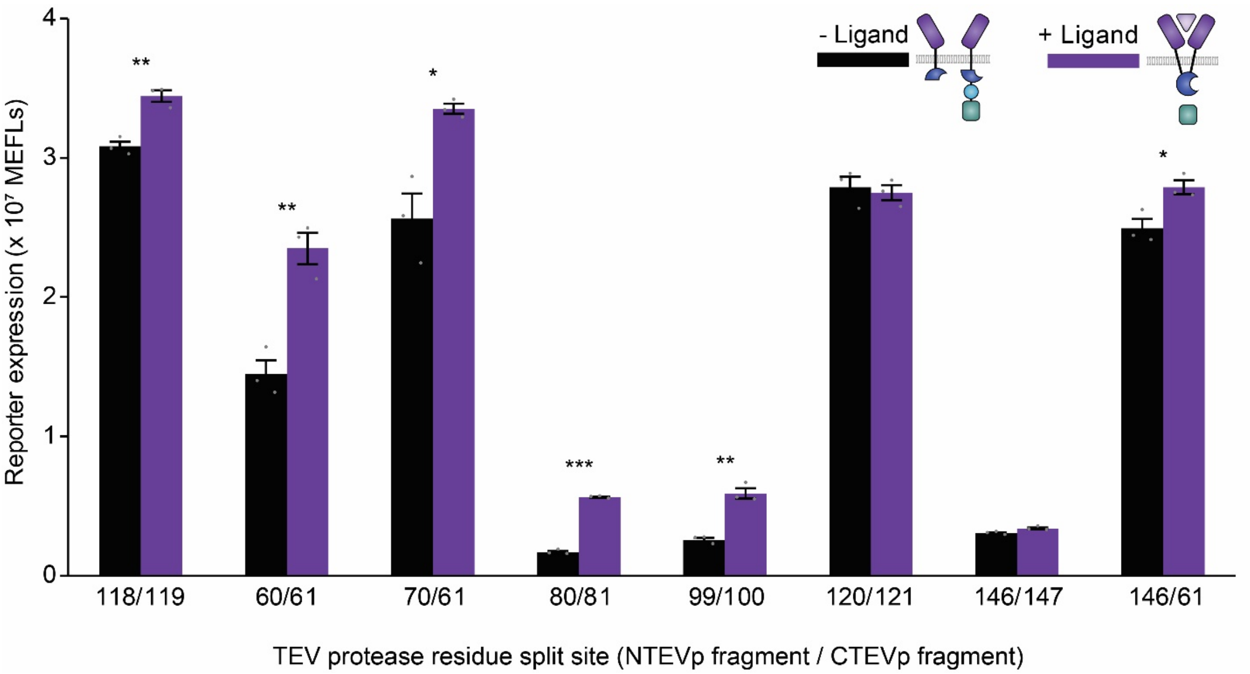


**Supplementary Fig. 2 Evaluation of alternative TEVp split sites.** Experimental analysis of TEVp splits other than 118/119 canonical split. Reporter expression driven by reconstituted TEVp cleavage of tTA measured with flow cytometry. Alternate splits were observed to have very high ligand independent signaling, or greatly diminished ligand independent and dependent signaling. This suggests that splits other than the canonical split do not have the optimal reconstitution propensity for the membrane bound environment. Error bars depict S.E.M. (*P ≤ 0.05, **P ≤ 0.01).

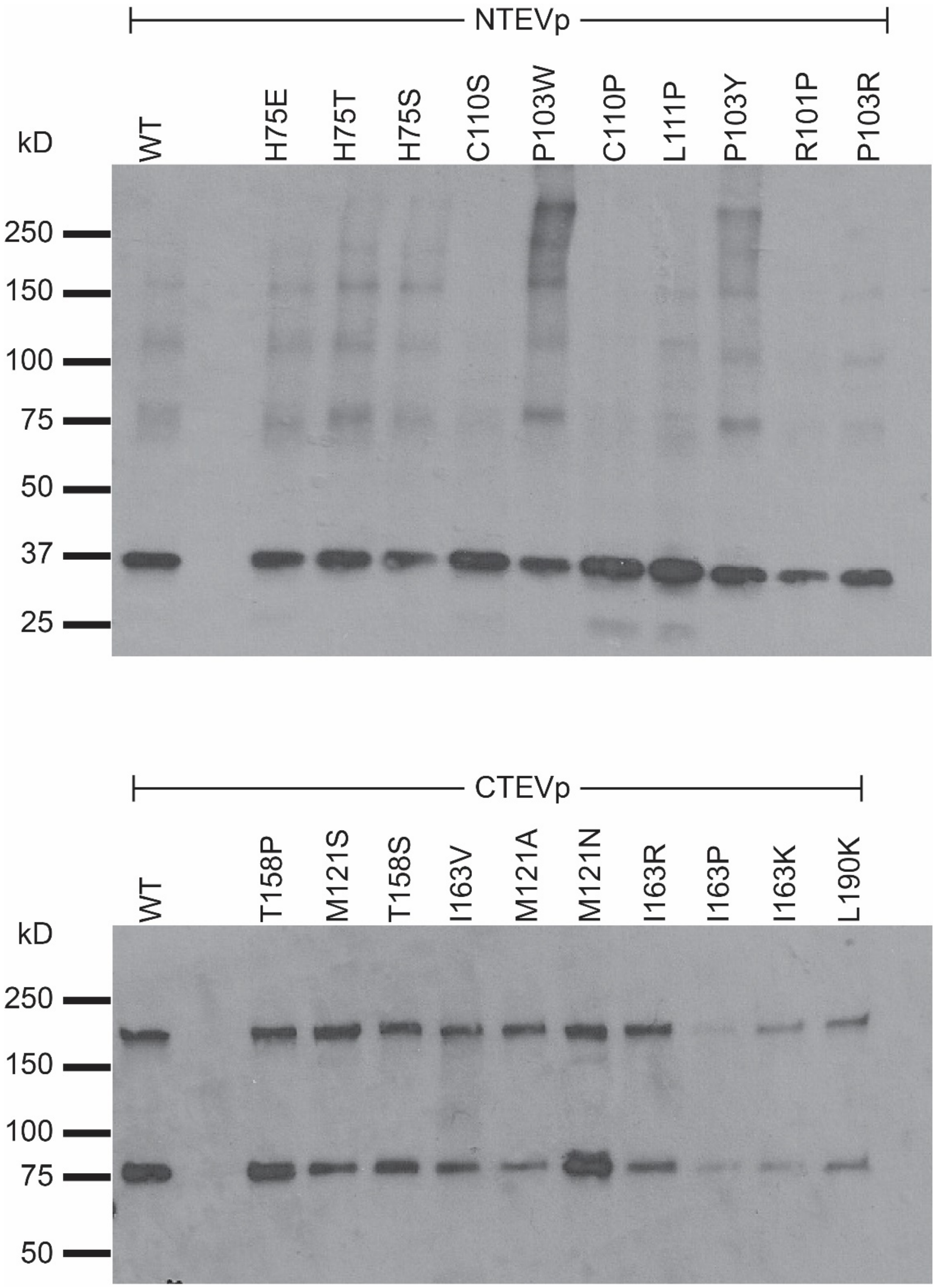


**Supplementary Fig. 3 Western blot analysis of mutant split TEVp expression levels.** Cells were transfected with single mutant chains which contain an N-terminal 3x-FLAG tag. Equal masses of protein (1 µg/lane) were loaded into each lane to investigate differences in chain expression level.

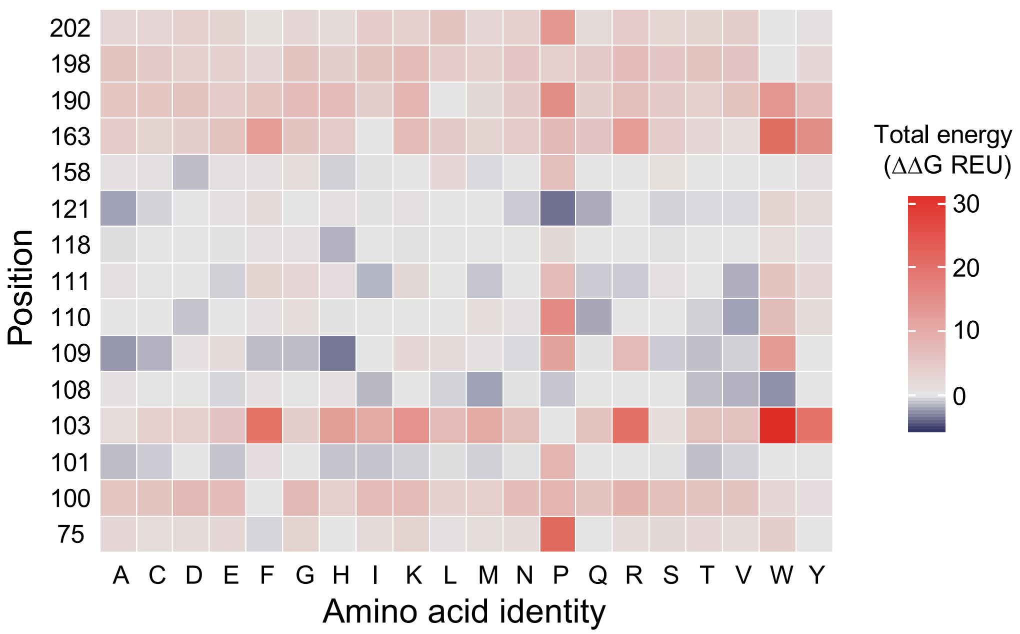


**Supplementary Fig. 4. Mutational scanning of high ΔSASA residues using Rosetta**. Energetic perturbations to total stability for all amino acid identities at each position are relative to the WT sequence.

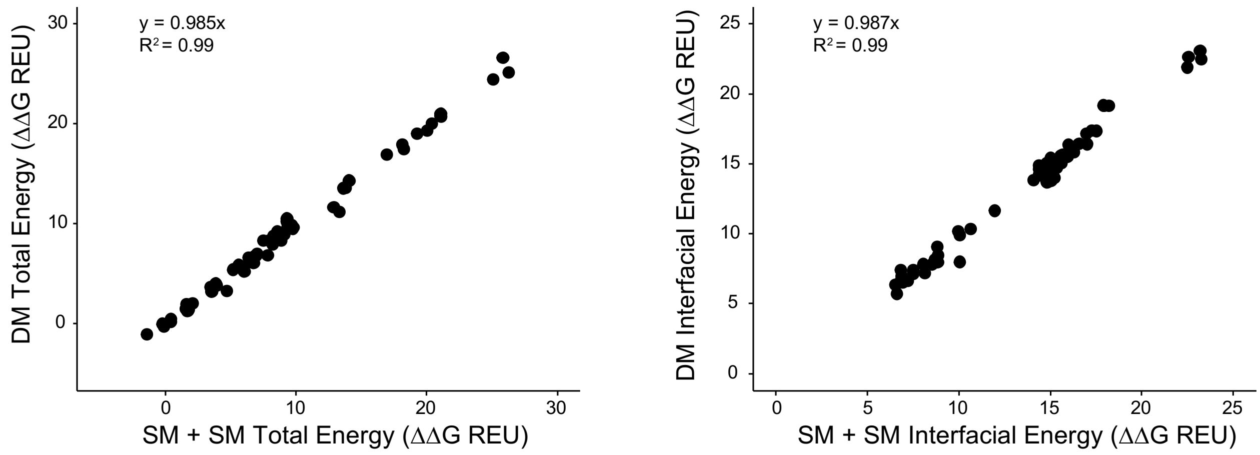


**Supplementary Fig. 5. Additivity of energetic perturbations for mutations as computed by Rosetta.** Data is shown for the sixty-seven double and paired mutants that can be created by combining the twenty single mutants evaluated in **Figure 2**.

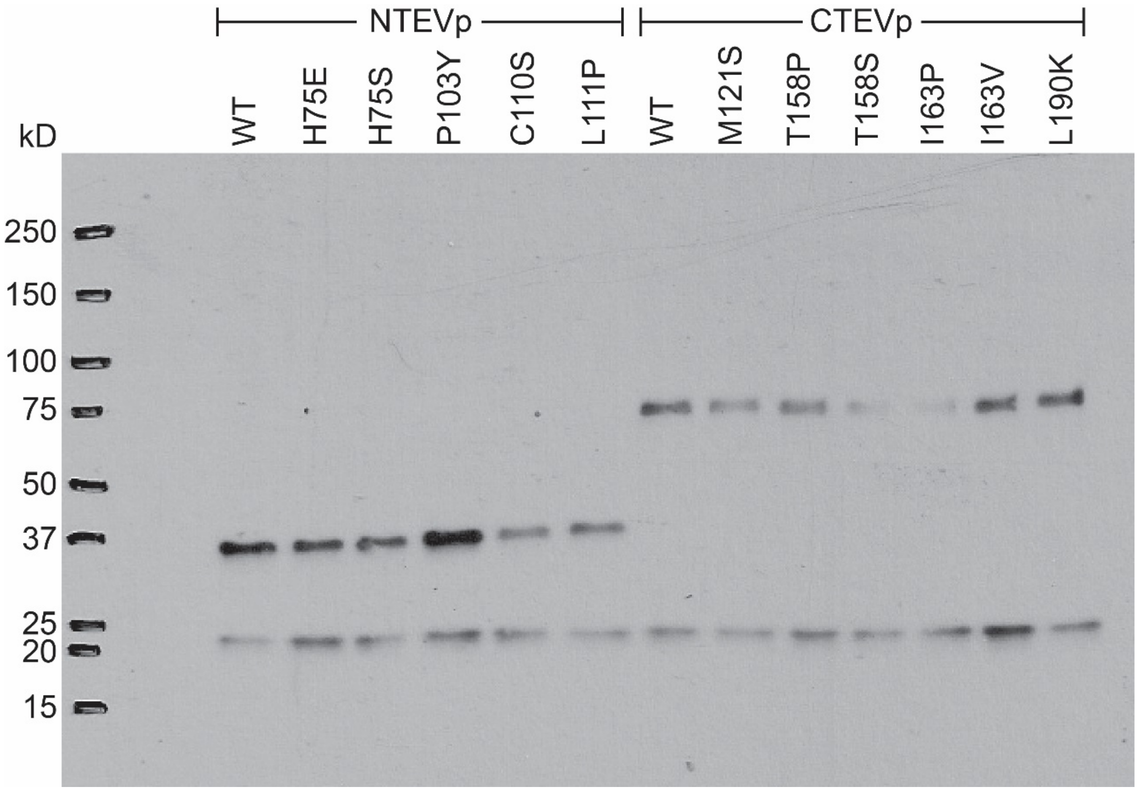


**Supplementary Fig. 6 Western blot analysis of mutant split TEVp expression levels.** Cells were transfected with single mutant chains which contain an N-terminal 3x-FLAG tag, and cells were co-transfected with a 3x-FLAG tagged NanoLuciferase as a normalization control (bottom band). Equal masses of protein (3 µg/lane) were loaded into each lane to investigate differences in chain expression level.

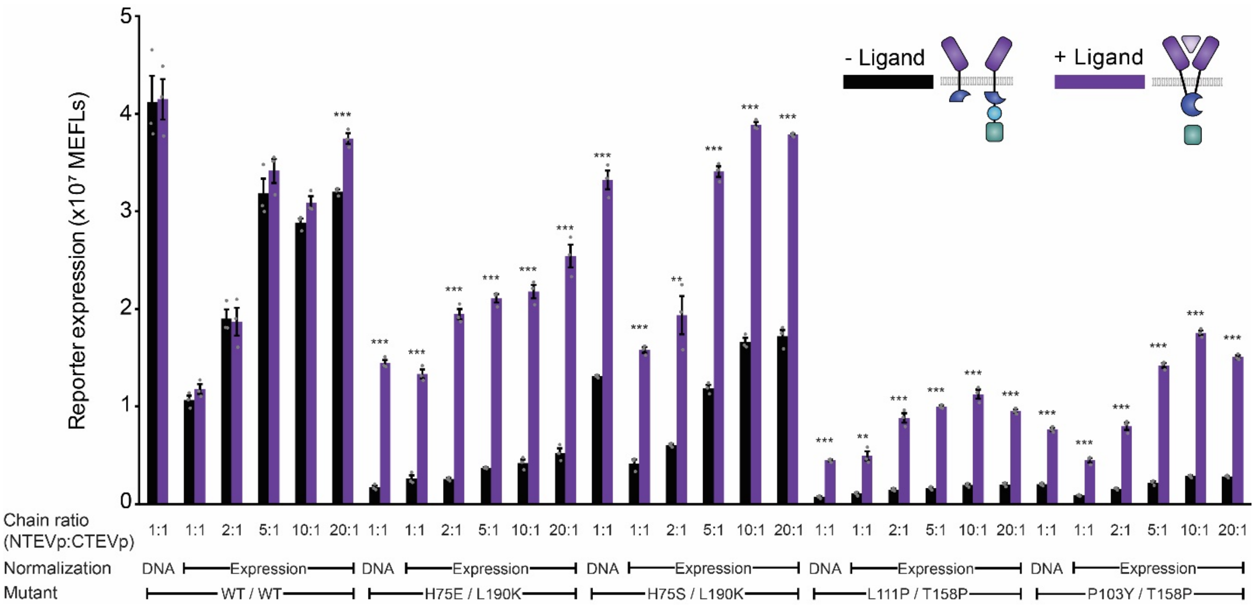


**Supplementary Fig. 7. Optimized splits yield robust performance across a wide range of expression levels**. Experimental analysis of DNA dose normalized and expression normalized select mutant pairs. Reconstituted TEVp cleavage of tTA measured by flow cytometry. Mutant pairs show inducible performance that is robust to the expression levels and ratios investigated, however WT performance is poor over this expression regime. Error bars depict S.E.M.

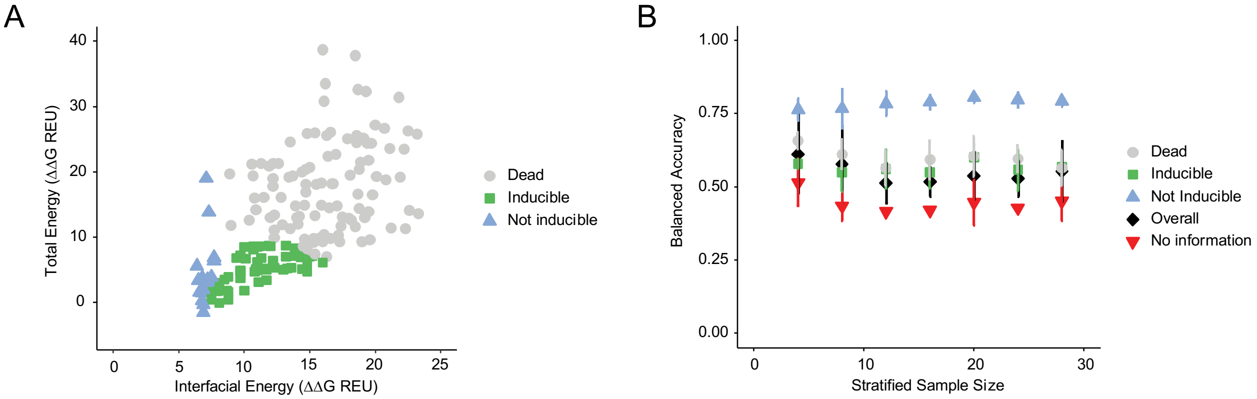


**Supplementary Fig. 8. Linear discriminant analysis (LDA) and retrospective bootstrapping**. **a,** Phenotype predictions for experimentally characterized variants (omitting the calibration set) using an LDA model trained on our actual 20-variant calibration set. Note that the phenotype boundaries determined by the LDA analysis are similar to those that were subjectively inferred (**Figure 2e**). **b**, Accuracy of phenotype predictions as a function of sample size using a retrospective bootstrapping approach of all 193 experimentally characterized variants. Classification accuracies were generated using LDA models trained from 10-fold stratified bootstrap partitioning of experimentally characterized variants into training sets**.** Accuracy was computed on variants not included in each training set. Error bars indicate the standard deviations across the 10-fold bootstrapping. Details for the analysis presented here are included in **Supplementary Note 2**.

**Supplementary Table 1.** List of vectors and plasmids used in this study. Plasmid maps are provided as **Supplementary Data 1**.

**Supplementary Note 1. Rosetta calculations used in this manuscript**

**Computational Interface Scanning**

Local Rosetta runs were performed with release version 3.9. The ref2015 scoring function was used for all calculations.

**Protein Preparation**

The structure of TEVp was obtained from the Research Crystallography for Structural Bioinformatics (RCSB) PDB (ID code: 1LVM). Chain designations A and B were used for residues 4-118 (NTEVp) and 119-221 (CTEVp), respectively. The structure was idealized, which sets bond lengths and angles to their ideal values and then minimizes the structure in the presence of coordinate constraints.

Commands:

```
${path_to_Rosetta}/main/source/bin/idealize_jd2.linuxgccrelease
        -database ${path_to_Rosetta}/main/database
        -in::file::fullatom
        -s 1LVM.pdb
        -no_optH false
        -flip_HNQ
```

**Interface Mutagenesis**

The idealized structure of TEVp was relaxed (100 iterations) using the default constrained relax script, and position constraints were added to backbone heavy atoms based on the crystal structure. During this relax procedure, Rosetta resfiles were used to incorporate single (or double) mutations.

Commands:

```
${path_to_Rosetta}/main/source/bin/relax.linuxgccrelease
        -database ${path_to_Rosetta}/main/database
        -relax::sequence_file always_constrained_relax.script
        -constrain_relax_to_start_coords
        -relax::coord_cst_width 0.25 -relax::coord_cst_stdev 0.25
        -s 1LVM.pdb
        -in::file::fullatom
        -no_optH false
        -flip_HNQ
        -nstruct 100
        -packing:resfile ${mutant}.resfile
        -relax:respect_resfile
```

*always_constrained_relax.script:*

```
repeat 5
    ramp_repack_min 0.02  0.01     1.0
    ramp_repack_min 0.250 0.01     1.0
    ramp_repack_min 0.550 0.01     1.0
    ramp_repack_min 1     0.00001  1.0
    accept_to_best
endrepeat
```

*example ${mutant}.resfile:*

```
AUTO
NATAA
start
75  A PIKAA A
```

**Interface Scoring**

The interfacial energy was computed for all relaxed structures of each variant using the rosetta_scripts application with the InterfaceAnalyzerMover. This mover calculates the total interaction energy between all residues in chain A (nTEV) with residues in chain B (cTEV).

Commands:

```
${path_to_Rosetta}/main/source/bin/rosetta_scripts.linuxgccrelease
    -database ${path_to_Rosetta}/main/database
    -parser:protocol interface_score.script
    -file:s ${mutant}.pdb
```

*interface_score.script*

```
<ROSETTASCRIPTS>
    <TASKOPERATIONS>
    </TASKOPERATIONS>
    <SCOREFXNS>
    </SCOREFXNS>
    <FILTERS>
    </FILTERS>
    <MOVERS>
```

```
		<InterfaceAnalyzerMover name="score_int" pack_separated="false" pack_input="false"
	packstat="false" interface_sc="true" ligandchain="B"/>
	</MOVERS>
	<PROTOCOLS>
	<Add mover_name="score_int"/>
	</PROTOCOLS>
</ROSETTASCRIPTS>
```

**Supplementary Note 2. Logistic regression analysis**

**Logistic regression**

We used logistic regression as a linear classification tool to relate experimentally determined phenotypes to the Rosetta computed energy landscape. During this type of analysis, hyperplanes in feature space are defined and used to separate observations belonging to different classes. All data preparation and logistic regression analyses were performed using R version 3.6.1. Stratified sampling and logistic regression using the linear discriminant analysis method (see commands below) were conducted with the dplyr and carat packages.

**Data stratification**

Stratified sampling reduces the likelihood of randomly sampling the same region of the energy landscape when selecting data to form calibration sets; sampling across the range of ΔΔG values increases the probability of sampling members of each phenotype, which is necessary for determining the energy partitions. The comprehensive Rosetta computed $\Delta\Delta G_{Total}$ vs $\Delta\Delta G_{Interfacial}$ energy landscape of all single (285) and double or paired mutants (37,905) was generated and divided into 4 evenly populated strata (I, II, III, and IV). To generate the strata, the data were first partitioned using the median $\Delta\Delta G_{Total}$, and then these subgroups were further partitioned by their median $\Delta\Delta G_{Interfacial}$. The strata boundaries were drawn from this ~38,000-member set because these would be the only data available at this stage if we were applying SPORT to a new split protein system. The 193 experimentally characterized TEVp variants were mapped to these strata for the retrospective stratified bootstrap sampling analysis.

<u>Commands:</u>

```
library(caret)

library(dplyr)

#LDA on our actual calibration set

cal <- read.csv("actual_calibration_set.csv", header=T)

new <- read.csv("stratified_after_calibration_set.csv", header=T)

train <- cal

test <- new

train_ddg <- cbind(train$ddG_total, train$ddG_interfacial)

train_pheno <- as.factor(as.character(train$phenotype))

test_ddg <- cbind(test$ddG_total, test$ddG_interfacial)

test_pheno <- factor(test$phenotype, levels=levels(train_pheno))

lda_model <- train(x=train_ddg, y=train_pheno, method="lda", metric="Accuracy")

lda_pred <- predict(lda_model, test_ddg)confusionMatrix(lda_pred, as.factor(test_pheno))


#Retrospective LDA using stratified bootstrap sampling of characterized variants

data <- read.csv("stratified_all_tested_data.csv", header=T)

for(size in 1:7) #since there are 4 strata, we are sampling 4, 8, …, 28 variants
```

```
{
    for(samp_set in 1:10)
    {                    set.seed(size*samp_set+100)
        strat_samp <- as.data.frame(data %>% group_by(strata) %>% sample_n(size)
%>% ungroup)
        train <- data[strat_samp[,1],]
        test <- data[-strat_samp[,1],]
        train_ddg <- cbind(train$ddG_total, train$ddG_interfacial)
        train_pheno <- as.factor(as.character(train$phenotype))
        test_ddg <- cbind(test$ddG_total, test$ddG_interfacial)
        test_pheno <- factor(test$phenotype, levels=levels(train_pheno))lda_model <-
    train(x=train_ddg, y=train_pheno, method="lda", metric="Accuracy")
        lda_pred <- predict(lda_model, test_ddg)
        cm <- confusionMatrix(lda_pred, as.factor(test_pheno))
        temp_byclass <- as.table(cm$byClass)
        temp_overall <- as.table(cm$overall)
        if(samp_set==1)
        {
            byclass <- temp_byclass
            overall <- temp_overall
        } else
        {
            byclass <- rbind(byclass, temp_byclass)
            overall <- rbind(overall, temp_overall)
        }
    }
    notInd <- subset(byclass, row.names(byclass)=="Class: not inducible")
    Ind <- subset(byclass, row.names(byclass)=="Class: inducible")
    Dead <- subset(byclass, row.names(byclass)=="Class: dead")
    all_overall <- overall[1:2,]
    row.names(all_overall) <- c('mean','sd')
    all_notInd <- byclass[1:2,]
```

```
row.names(all_notInd) <- c('mean','sd')
all_Ind <- all_notInd
all_Dead <- all_Ind
for(param in 1:length(overall[1,]))
{
	all_overall[1,param] <- mean(overall[1:length(overall[,1]),param], na.rm = TRUE)
	all_overall[2,param] <- sd(overall[1:length(overall[,1]),param], na.rm = TRUE)
}
for(param in 1:length(byclass[1,]))
{
	all_notInd[1,param] <- mean(notInd[1:length(notInd[,1]),param], na.rm = TRUE)
	all_notInd[2,param] <- sd(notInd[1:length(notInd[,1]),param], na.rm = TRUE)
	all_Ind[1,param] <- mean(Ind[1:length(Ind[,1]),param], na.rm = TRUE)
	all_Ind[2,param] <- sd(Ind[1:length(Ind[,1]),param], na.rm = TRUE)
	all_Dead[1,param] <- mean(Dead[1:length(Dead[,1]),param], na.rm = TRUE)
	all_Dead[2,param] <- sd(Dead[1:length(Dead[,1]),param], na.rm = TRUE)
}
if(size==1)
{
	mean_overall <- all_overall[1,]
	sd_overall <- all_overall[2,]
	mean_notInd <- all_notInd[1,]
	sd_notInd <- all_notInd[2,]
	mean_Ind <- all_Ind[1,]
	sd_Ind <- all_Ind[2,]
	mean_Dead <- all_Dead[1,]
	sd_Dead <- all_Dead[2,]
} else
{
	mean_overall <- rbind(mean_overall,all_overall[1,])
	sd_overall <- rbind(sd_overall,all_overall[2,])
	mean_notInd <- rbind(mean_notInd,all_notInd[1,])
```

```
                    sd_notInd <- rbind(sd_notInd,all_notInd[2,])
                    mean_Ind <- rbind(mean_Ind,all_Ind[1,])
                    sd_Ind <- rbind(sd_Ind,all_Ind[2,])
                    mean_Dead <- rbind(mean_Dead,all_Dead[1,])
                    sd_Dead <- rbind(sd_Dead,all_Dead[2,])
          }
}
```
